# Lessons learned from applying eDNA surveying to diadromous fish detection across the north-east Atlantic region

**DOI:** 10.1101/2025.01.31.635873

**Authors:** Mukesh Bhendarkar, Cristina Claver, Iñaki Mendibil, Natalia Fraija-Fernández, David J. Nachón, Phil I. Davison, Tea Bašić, Ciara O’Leary, William K Roche, Iker Azpiroz, Marie-Laure Acolas, Aitor Lekuona, José Ardaiz, Estibaliz Diaz, Patrick Lambert, Geraldine Lassalle, Naiara Rodriguez-Ezpeleta

## Abstract

Regular monitoring of diadromous fishes is critical to inform their management and conservation. Yet, the *in-situ* data collection these species is challenging due to their complex life cycle and low abundance. Focusing on the sea lamprey (*Petromyzon marinus*, Petromyzontidae) and the European shads (*Alosa alosa* and *A. fallax*, Clupeidae), emblematic diadromous fishes in the Northeast Atlantic region, this study leverages the use of water environmental DNA (eDNA) samples to monitor their distribution range. For that aim, we developed quantitative PCR (qPCR) and digital PCR (dPCR) assays and applied them to detect sea lamprey and European shad DNA in a network of 44 river basins across Spain, France, Ireland, and the UK. We found that qPCR efficiently detected presence/absence of shads, while the higher sensitivity of dPCR was essential for detecting the lower abundant and partly sessile behaving sea lamprey in the amount of water collected. Moreover, sea lamprey showed significantly lower eDNA copies per litre of water compared to shads, probably due to their larvae spending several years burrowed within soft sediments, reducing eDNA shedding into the water column. The integration of historical datasets with this snapshot wide-ranging study enhances our understanding of the distribution of sea lamprey and European shad in Atlantic rivers. Importantly, the lessons learned within this international collaboration are critical towards a prevailing framework for conservation of migratory fishes, highlighting the need of well-designed sampling strategies coupled with species-specific assays applied to eDNA samples to bust long-term monitoring efforts of diadromous species.

## Introduction

Anthropogenic activities, together with climate change, can significantly modify natural habitats and ecosystems (Chu et al., 2005; Dodds et al., 2013). This is particularly critical for diadromous species, whose complex life cycles, involving a variety of habitats between rivers and open ocean, make them more vulnerable to any alterations (Chaparro-Pedraza & de Roos, 2019; Limburg & Waldman, 2009; Tamario et al., 2019). Diadromous fishes are emblematic species with crucial ecological roles, providing numerous ecosystem services (Almeida et al., 2023; Ashley et al., 2023; Naiman et al., 2002). Thay can either be catadromous (migrating to sea to spawn) or anadromous (migrating to rivers to spawn). The sea lamprey (*Petromyzon marinus* Linnaeus, 1758) and European shads (*Alosa alosa* Linnaeus, 1758 and *A. fallax* Lacépède, 1803) are ecologically, evolutionarily, and economically important anadromous species, which are present along the eastern Atlantic coast (Almeida P R & Rochard, 2015; Wilson & Veneranta, 2019). Their distributions and abundances have decreased over time due to the synergistic effect of anthropogenic impacts, including climate change, so that their core distributions are now restricted to southwestern Europe mostly (Almeida et al., 2021; Lassalle et al., 2008; Nachón et al., 2020). Despite being categorized by the International Union for Conservation of Nature (IUCN) as “Least Concern” (LC) globally and at the European level (Freyhof, 2010a; Freyhof, 2010b; Freyhof & Kottelat, 2008a, 2008b; NatureServe, 2013), these species are listed as “Threatened” (T) or declining in several European countries (Almeida P R & Rochard, 2015; Limburg & Waldman, 2009; OSPAR Commission, 2009a, 2009b). For instance, in France and Great Britain, *A. alosa* is listed as “Critically Endangered” (CR), while *A. fallax* is categorized as “Near Threatened” (NT) in France and “Vulnerable” (VU) in Great Britain. Additionally*, P. marinus* is classified as “Endangered” (EN) in France and NT in Ireland (King et al., 2011; Nunn et al., 2023; UICN Comité français, 2019). This high level of conservation concern at national levels and across boundary distribution of the species calls for concerted inter-basin conservation and management efforts (Guo et al., 2016; ICES, 2003; Kritzer et al., 2022; Ouellet et al., 2022).

In this context of generalized decline, obtaining accurate information about diadromous species occurrence is essential to identify key periods and habitats when and where feeding, breeding and migration occur. Continual change, especially alterations in species home river ranges due to various pressures, emphasizes the importance of understanding their movements and habitats (Beaulaton et al., 2008; ICES, 2003). Yet, characterizing the distribution of fish spatially and temporally is a challenging task, especially when considering factors like the scale of distribution, the complex life history of diadromous species, and resource constraints (Ciannelli et al., 2008). Conventional methods, such as mark recapture and electrofishing have been routinely used, but have their own challenges, including difficulties in deployment in all habitats (Lapointe et al., 2006; Pont et al., 2021). Moreover, these methods are not only intrusive but can also be costly and labour-intense (Lapointe et al., 2006; Pont et al., 2021), as well as inefficient in detecting low abundant species (Cantera et al., 2019; Piggott et al., 2021). Therefore, using a cost-effective, scalable and non-invasive sampling method that increases the probability of detection is key for monitoring rare aquatic species with such a restricted temporal and wide spatial distribution as anadromous fishes (Westhoff et al., 2022). The analysis of environmental DNA (eDNA) is a promising and rapidly evolving approach for aquatic species distribution monitoring (Bhendarkar & Rodriguez-Ezpeleta, 2024; Nagarajan et al., 2022; Pawlowski et al., 2021; Rodríguez-Ezpeleta et al., 2021), with potential adaptability in a changing world (Thomsen et al., 2024). Traces of organisms in the form of gametes, faeces, skin cells, saliva, blood, and other bodily substances contain eDNA (Bohmann et al., 2014) that can be utilized to determine the presence of a species within a specific area (Abbott et al., 2021; Davison et al., 2019). This methodology has been instrumental in examining spatio-temporal patterns around spawning habitats in rivers for *P. marinus* (Bracken et al., 2019; Moser et al., 2021) and *Alosa* spp. (Antognazza et al., 2021; Antognazza et al., 2019). Moreover, eDNA-based monitoring has made it possible to identify prospective nursery grounds and distribution of sea lamprey larvae (Baltazar-Soares et al., 2022). Conversely, the eDNA-based approach is also used to assess surveillance and control measures for invasive populations of sea lamprey in other parts of the globe where the species is considered as a pest (Gingera et al., 2016; Schloesser, 2018; Tkachuk & Dunn, 2020).

In this study, we conducted a broad-scale survey using eDNA to monitor the geographical distribution of sea lamprey and European shads in their known range of the North-East Atlantic region. By using eDNA-based surveys, the goal was to provide a non-invasive, efficient and scalable approach to monitor these species across multiple river basins in Spain, France, Ireland and the UK. We aimed to address four objectives: i) to assess the regional distribution of sea lamprey and European shads; ii) to evaluate the efficacy of eDNA analysis as a biomonitoring tool within this context; iii) to compare the variability of eDNA detections between quantitative PCR (qPCR) and digital PCR (dPCR) methods; iv) to determine freshwater migration limits of sea lamprey and shads within river-estuary systems during their upstream migration. The first two objectives were achieved by comparing qPCR-based eDNA detections with evidence-based knowledge of species occurrence across 44 river basins in Spain, France, Ireland and the UK, while the third and fourth objectives were based on dPCR-based eDNA detections from river estuaries within Spain. As methodology continues to evolve, the integration of eDNA-based monitoring is likely to become increasingly essential in the assessment of diadromous species. This work improves our understanding of the practicality and efficiency of eDNA analysis at a river basin scale, providing a robust framework for future research and highlighting the importance of a multi-approach to eDNA analysis interpretation.

## Material and Methods

### Sampling location and reference data set

Water samples were collected from a network of 44 river basins across Spain (20 rivers; 168 samples), France (2 rivers; 9 samples), the UK (15 rivers; 53 samples), and Ireland (7 rivers; 68 samples – including 3 at sea close to the river mouth). In the Basque region (Spain), additional water samples were taken from four estuaries: Bidasoa, Oiartzun, Urola and Deba (15 samples) to investigate the abundance and distribution of eDNA along the stretch of river leading to the upstream sites through digital PCR. The sampling occurred over different years (2019 to 2021) in different regions, resulting in a total of 313 samples (**Figure 1; Table S1**). The selection of sampling sites was based on the known historic distribution of sea lamprey and shads in their native range within these basins coupled with opportunistic sampling (**Table S1**). Further details on transboundary eDNA sampling and analytical methods can be found in the DIADES Project manual (https://diades.eu/wp-content/uploads/2020/04/WP6_1_Manual_Final_Version-compress%C3%A9.pdf).

**Figure 1.**
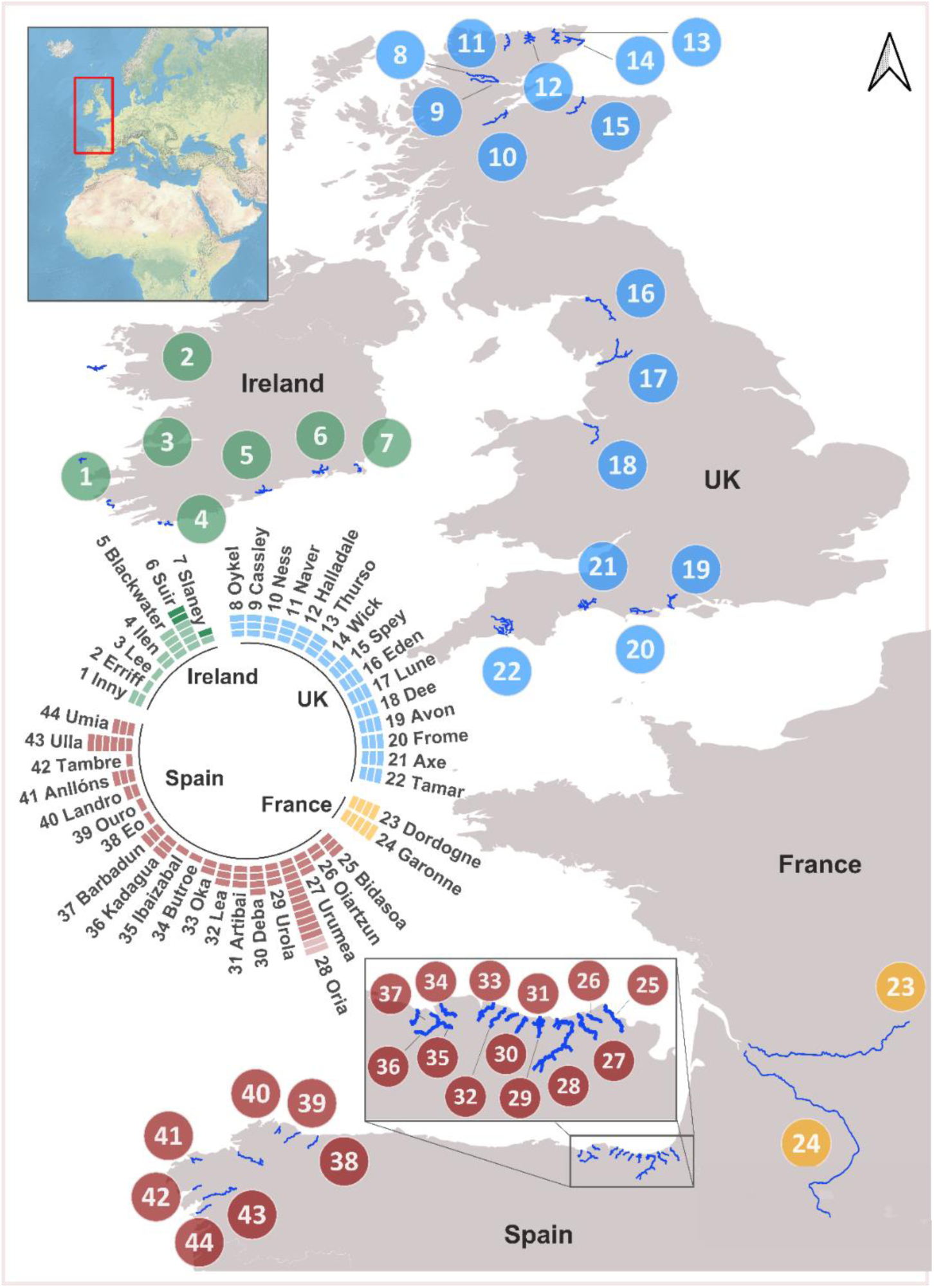
Geographical location of river basins sampled for in this study; main stems are represented with a blue line. Each section of the bar charts indicates a sampling point analysed through qPCR. Darker sections indicate sea samples taken near the river mouth (Slaney and Suir rivers) and lighter sections indicate sampling points located in affluents of the main river basin (Oria river). See Table S1 for additional details.

To place eDNA-based findings in the context of expected presence or absence, a historical evidence reference database for the sea lamprey and shads for each eDNA sampling site was created by combining presence/absence data from electrofishing, traps, and net operations, as well as presence of downstream barriers; evidence has been sourced from the scientific literature or from database repositories (**Table S1**). For each species, each eDNA sampling point was categorized as “Expected” where there was evidence of historical presence; “Not expected” where there was evidence of absence or an impassable downstream barrier; and “Unknown” where there was no evidence to indicate the presence or absence of the species.

### Water sample collection and DNA extraction

The volume of water sample filtered ranged from 1 to 30 litres (**Table S2**). Spanish and French samples were filtered through 0.45 μm pore size Sterivex filters and VigiDNA® crossflow filtration capsules, respectively. Irish samples filtered through 1.50 μm pore size Whatman filters, while for the UK samples, 0.22 μm pore size Sterivex filters were utilized (**Table S2**). All filters were kept frozen until further processing. For DNA extraction, the Qiagen QIAamp DNA kit was used for Spanish and Irish samples, while the French samples were processed using an in-house procedure (Pont et al., 2018), and UK samples were extracted using the DNeasy Power Water Sterivex kit (Qiagen) following the manufactureŕs instructions. Sample handling and pipetting were meticulously performed within a specialized hood to prevent contamination. Each filter was extracted and analyzed individually, ensuring the integrity of the DNA extracts from each separate water samples. The extracted DNA was then transported to AZTI in Spain for further analysis. The concentration of the extracted DNA was quantified using UV spectrometry (Thermo Scientific NanoDrop™ ND-1000) and Qubit^TM^ fluorometer (Life Technologies), and its quality was assessed through agarose gel electrophoresis.

### Species-specific detection assay development

To detect both species of European shads inhabiting European waters (*A. alosa* and *A. fallax*), we designed genus-specific primers and a probe for *Alosa* spp. targeting a 94bp fragment of the mitochondrial cytochrome b (*cytb*) gene. To do so, all *Alosa cytb* available sequences were retrieved from GenBank (https://www.ncbi.nlm.nih.gov). These sequences were aligned and a suitable region in which to develop the specific assay was identified using the BioEdit software (Kirmani, 2015). To ensure the specificity of the primer and probe sets, an *in silico* analysis was performed using BLAST (Altschul SF, 1990), and no potential amplification of non-target DNA was detected. For the detection of *P. marinus*, we utilized the assay developed by Moser et al. (2021). The sequences of the primers and probes used in this study are detailed in **Table 1**.

**Table 1.**
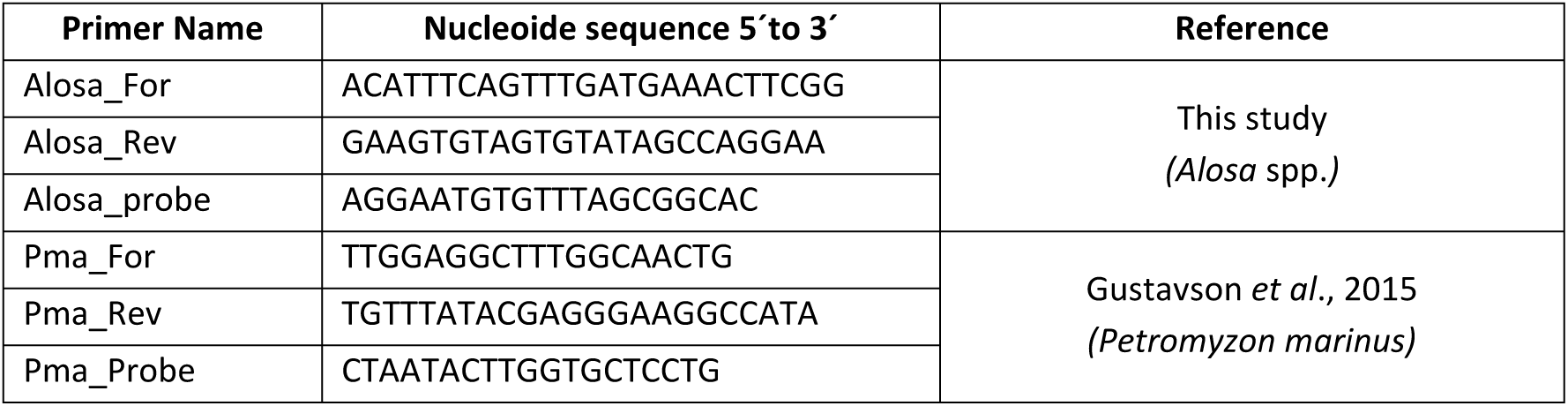
Primer and probe sequences used for European shads and sea lamprey specific amplification and detection.

### Quantitative PCR (qPCR) analysis

Quantitative PCR analyses were performed by combining the 5 µl of TaqMan™ Fast Advanced Master Mix (Applied Biosystems), 0.2µl of each primer (forward and reverse) and probe (10 µM), 2 µl extracted DNA (10 ng) and 2.4 µl Milli-Q water, to adjust the total reaction volume to 10 µl. The procedure was carried out independently to detect the European shads and sea lamprey under the following conditions: for *Alosa* spp. (hereafter referred to as shads), initial 50°C for 2 min and 95°C for 10 min, followed by 50 cycles alternating between 95°C for 3 s and 60°C for 20 s; and for *P. marinus* (sea lamprey), 50 cycles alternating between 95°C for 3 s and 55°C for 20 s. A total of five PCR replicates were analyzed for each sample, along with positive and negative controls. Genomic DNA (gDNA) extracted from tissue samples of specimens collected in 2019 from the river basins in the Basque Country by EKOLUR Environmental Consulting (Spain) was used as a positive control, with a 10-fold standard dilution series included on each qPCR plate. A sample was considered positive for the species if at least one qPCR replicate yielded a cycle threshold (Ct-value, indicating the number of cycles in qPCR needed to detect the DNA signal) value less than 40 (Takahara et al., 2020; Westhoff et al., 2022). To validate that the PCR was not inhibited, an additional analysis adding a known amount of DNA (0.1 ng) of a targeted species (Internal Positive Control - IPC) to each sample was performed. The composition of the PCR mixture as well as the thermocycling conditions were the same as above, but the reaction included 2 µl of IPC, with an adjustment to the volume of water. A sample was identified as inhibited if the Ct value was delayed or failed to amplify in this PCR.

### Digital PCR (dPCR) analysis

Digital PCR (dPCR), utilizing the QIAcuity One, 5plex (Qiagen), was used to validate and compare the detection sensitivities between qPCR and dPCR on samples collected from Spanish rivers. Additionally, we analyzed a new set of samples that were collected along transects extending from the estuaries of rivers Bidasoa, Oiartzun, Urola and Deba. The primer and probe sets utilized in the dPCR assays were consistent with those used in the qPCR analyses, with ROX-labelled probes designated for shad detection and Cy5-labelled probes for sea lamprey detection. Each dPCR reaction contained 10 µl of QIAcuity Probe PCR Kit, 3.2 µl of probe mix, 2 µl of extracted DNA, and 24.8 µl of Milli-Q water, resulting in a total reaction volume of 40 µl. This 40 µl reaction volume was then transferred into microfluidic dPCR nanoplates (QIAcuity) capable of accommodating 24 samples, with each well divided into up to 26,000 partitions. Each of these partitions enables individual PCR reactions to occur within them. The nanoplate was then loaded onto the digital PCR instrument (QIAcuity One, 5plex) and subjected to an automated workflow after quantifying the cycling protocol. The thermal cycle was executed under the following conditions: an initial step at 95°C for 2 minutes, followed by 40 cycles alternating between 95°C for 15 seconds and 60°C for 60 seconds. Each plate included positive controls (standard dilution series with tissue samples) and negative controls (no template controls, NTC). For detection, the ROX fluorescence channel was engaged to monitor shad amplification, whereas the Cy5 fluorescence channel was used for sea lamprey amplification. Fluorescent images from all PCR wells were acquired and analyzed; partitions indicating the presence of the target molecule were discerned by their elevated fluorescence intensity.

The common threshold value of fluorescent intensity (RFU) was set manually, in accordance with Qiagen handbook recommendations (Qiagen, 2022), with both negative and positive controls in each reaction (Passera et al., 2023). This choice allowed us to clearly distinguish between positive and negative partition clouds. The blank negative control consistently showed negative results for DNA copies. The eDNA copy number per reaction volume were calculated by QIAcuity Software Suite using Poisson statistics on the ratio of positive and negative partition (Qiagen, 2022).

### Quantification of dPCR and qPCR analysis

To determine the concentration of sea lamprey and shad eDNA in water samples, in copies per litre (copies/L), we started by assessing the concentration of the DNA template within the reaction volume, which was measured in copies per microlitre (copies/µL). Subsequently, we quantified the concentration of the DNA in the extraction eluate, which was based on a standard volume of 100 µL. For each individual water sample, we used the following formula to compute the overall eDNA concentration:

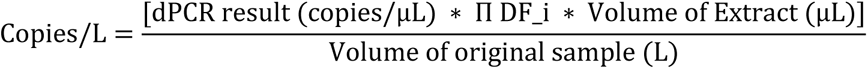

Where, Π DF_i is the product of all dilution factors in sample.

We conducted a linear regression analysis, utilizing the R software (R Core Team, 2021), to examine the relationship between the eDNA concentrations obtained from dPCR assays (expressed in copies/L) and the CT values via qPCR.

### Confusion matrix analysis

A confusion matrix is a popular measure in machine learning that sums up the performance of a classification model into a table (Sokolova & Lapalme, 2009). It works for binary and multi-class classification. We constructed two separate confusion matrices. The first matrix was based on the detected/not detected data obtained from the qPCR-based eDNA assays and compared to the expected/not expected data derived from reference databases (**Table S1**). This allowed us to assess the accuracy of eDNA method for detecting sea lamprey and shads compared to their known expectations. This matrix serves to quantify how well our eDNA analysis matches the expected presence or absence of the species. Samples categorized as ‘Unknown’ were excluded from the analysis. The second confusion matrix compared the detection rates obtained from qPCR assays and dPCR assays, allowing a direct comparison of these two molecular techniques in their performance in detecting sea lamprey and shads. The resulting confusion matrix was used to calculate different performance metrics such as accuracy (the proportion of correct predictions both true positives and true negatives out of all predictions), sensitivity (correctly identify true positives), and specificity (correctly identify true negatives). These metrics calculated as:

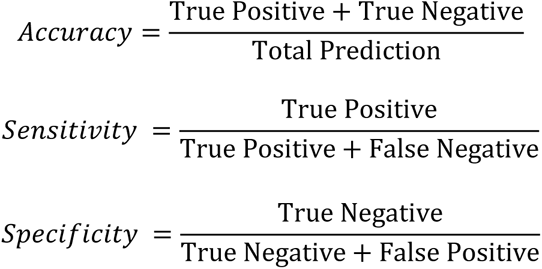

The confusion matrix analysis was performed using the R software (R Core Team, 2021).

## Results

### Comparative performance of eDNA analysis

The comparison between the evidence-based historical records of the existence of the species and the current eDNA detection outcomes revealed that among the 58 sampling sites where the presence of sea lamprey was expected, eDNA detection occurred only at 17 sites. No sea lamprey eDNA was detected in the 37 unknown and 38 not expected sites (**Figure 2a; Table S3**). For shads, eDNA detection was successful at 14 out of the 35 sites where the species was historically anticipated. In addition, positive detections were recorded at 4 of the 31 sites that were previously ‘unknown’ and at 3 of the 67 sites where the species was not expected based on the reference dataset (**Figure 2b; Table S3**).

**Figure 2.**
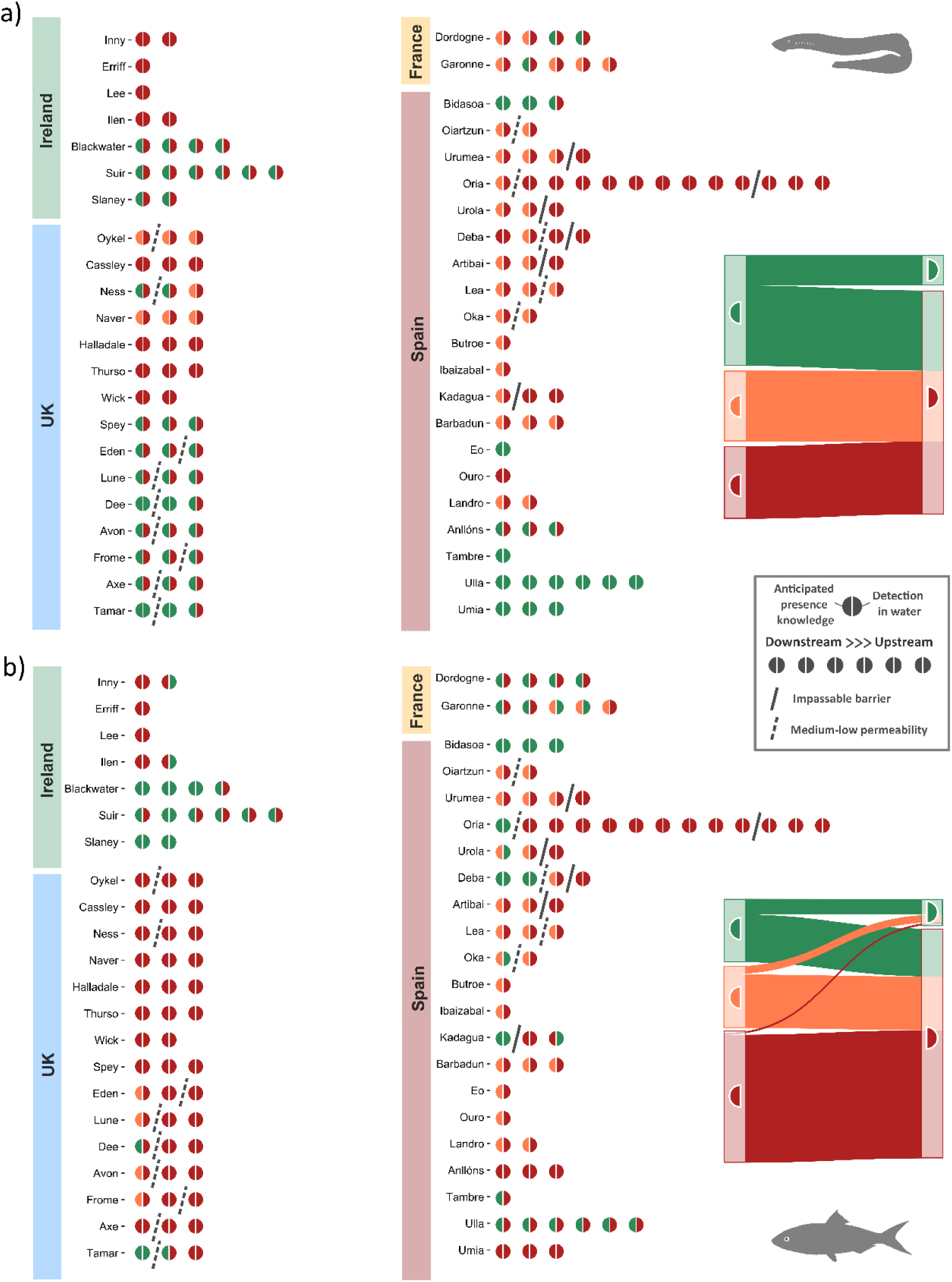
Detections of a) sea lamprey and b) European shads in European rivers using qPCR. Each dot represents one sampling point in the river basin, from downstream (left) to upstream (right). Straight lines indicate unpassable barriers and dashed lines, barriers with medium permeability. Left side of dots indicate if species presence (green) or absence (red) is expected or is unknown (orange) based on reference data (Table S1), whereas right side dots indicate detection (green) or non-detection (red) of the species using eDNA. Additionally, the Sankey diagrams on the right illustrate overall flow between expected species presence/absence (left) and their corresponding eDNA detection outcomes (right).

The confusion matrix output (**Figure 3**), excluding the ‘unknown’ category, offers a quantitative overview of the of eDNA detection (qPCR method) against anticipated presence/absence derived from reference data sources. For sea lamprey, eDNA detection exhibited an overall accuracy of 57.29% and a sensitivity of 29.31%, indicating that eDNA analyses using qPCR successfully detected sea lampreys in only a small fraction of the sites where they were expected. The specificity of 100% shows that in all the sites where reference data predicted sea lamprey absence, the qPCR analyses concurred (**Figure 3a**). For shads, eDNA detection using qPCR showcased an accuracy of 76.47% and a sensitivity of 40.0%, emphasizing its proficiency in confirming the presence of shads in locations where it was anticipated according to reference data. Meanwhile, a specificity of 95.52% indicates that in places where reference data predicted the absence of shads, eDNA detection often supported this absence (**Figure 3b**).

**Figure 3.**
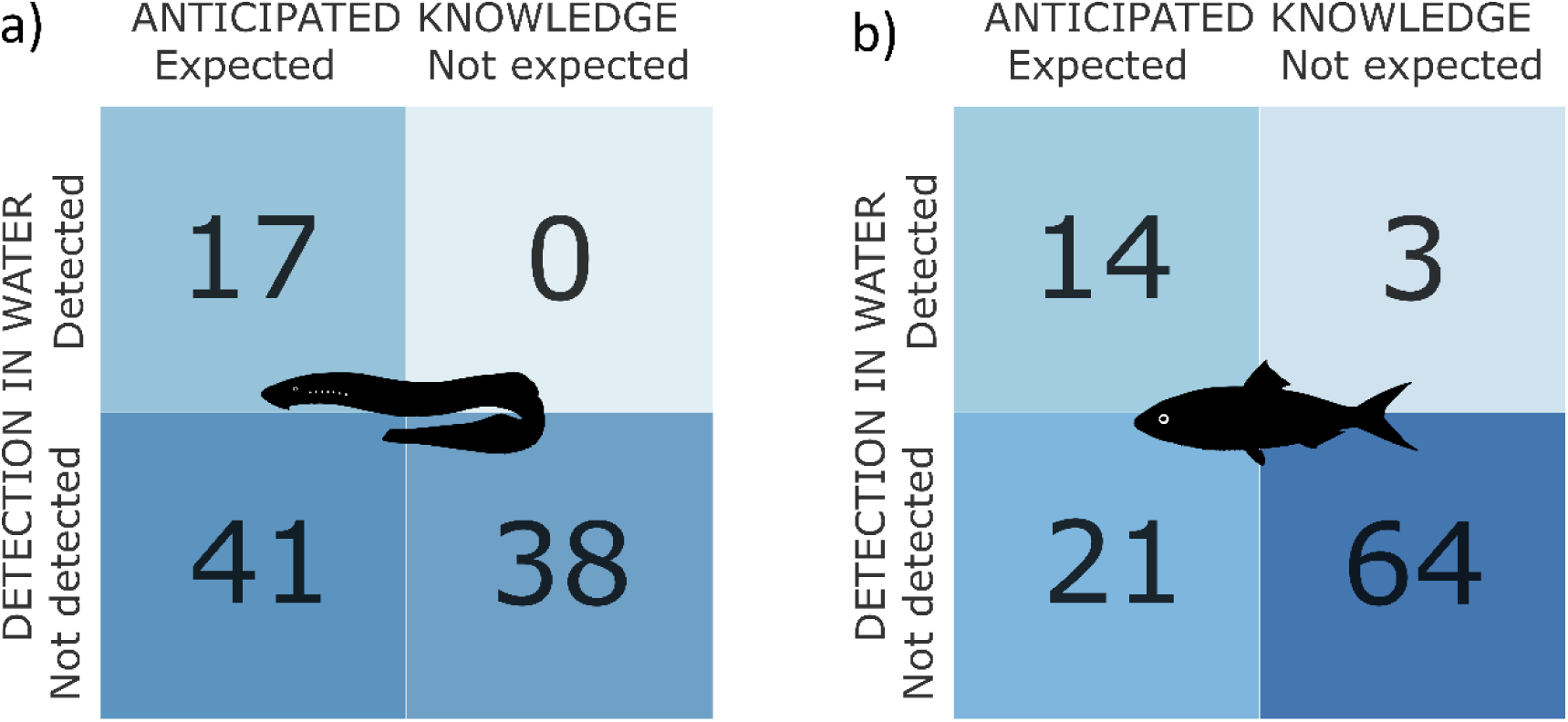
Integrative analysis of expected species presence/absence and eDNA detection in the form of a confusion matrix for a) sea lamprey and b) European shads, where the number of sites for each combination of reference data vs eDNA based result (excluding sites for which the species presence/absence was unknown) is indicated.

### River specific detection rate

Sea lamprey presence was confirmed exclusively in the Bidasoa, Eo, Tambre, Ulla, Umia rivers in Spain, and the Welsh Dee and Tamar rivers in the UK (**Figure 2a**). Despite evidence of sea lamprey presence in French, Irish, and other UK rivers, the species was not detected with eDNA. In all sites where their presence was unexpected, the qPCR assay provides support for their absence. Conversely, presence of shads was confirmed in the Garonne River in France, the Bidasoa River, downstream of the Oria, Deba, and Kadagua rivers in Spain, the Slaney, Suir, Blackwater, Ilen, and Inny rivers in Ireland, and the Tamar River in the UK (**Figure 2b**). Notably, upstream sites along the rivers Inny, Ilen in Ireland, and Kadagua in the Basque country, where the presence of the species was not expected, resulted in positive detection. Yet, it should be noted that only one replication (over five PCR) at each of these rivers resulted in a positive detection with low detection (CT>36). Additionally, positive detections were recorded along the rivers Garonne in France, Urola, and Oka in the Basque Country, which were previously classified as ‘unknown’. In the Basque country, specifically Bidasoa, Oiartzun, Urumea, Oria, Urola, and Deba, where we conducted eDNA sampling for two consecutive years (2019 and 2020), results showed a consistent detection patterns observed for sea lampreys and shads over the course of both years, reinforcing the utility and the reliability of our assays in monitoring these species.

### Insight from digital PCR (dPCR) and quantitative PCR (qPCR) assessment

While qPCR was used for all samples from Spain, UK, Ireland, and France, a more sensitive dPCR assay to allow more detailed interpretation was applied to the Spanish samples, from which enough DNA was available for additional analyses. Comparisons between dPCR and qPCR results showed similar detections rates in most sampling sites, except in Basque rivers where sea lamprey was detected by dPCR but not by qPCR (**Figure 4a; Table S3**). However, negative qPCR results for sea lamprey in most samples posed a challenge for holistic comparisons. It is important to note that qPCR analysis involved five replicates for each sample, and the CT value represents the average of the positive values, whereas dPCR analyses were conducted with single replicate, but whose result is the compilation of thousands of partitions (i.e., 26 thousand PCR independent reactions) within the reaction. The confusion matrix also reflects that dPCR was able to be detect 23 sites where qPCR failed, demonstrating an accuracy of 80.4%, a sensitivity of 50.0% and a specificity of 83.8% (**Figure 4b**). These metrics indicate that dPCR has a higher detection capability than qPCR, particularly for sea lamprey. For shads, the linear regression between mean qPCR CT values and log-transformed copies/L determined by dPCR underscores a positive correlation of 0.88 **(Figure 4a**). This suggests that an increase in eDNA copies corresponds to an enhanced detection rate through qPCR CT values and replications. Most sampling sites displayed a consistent detection rate compared to qPCR, indicating uniformity in the findings across different locations. In all Galician rivers, shad remained undetected, consistent with the qPCR results. In four sites dPCR did not yield positive while qPCR did, while with high CT values. The confusion matrix for shads showed that the accuracy of dPCR was 96.2%, the sensitivity 92.5%, specificity of 98.10% demonstrating a high level of consistency with the qPCR detection rate (**Figure 4b**).

**Figure 4.**
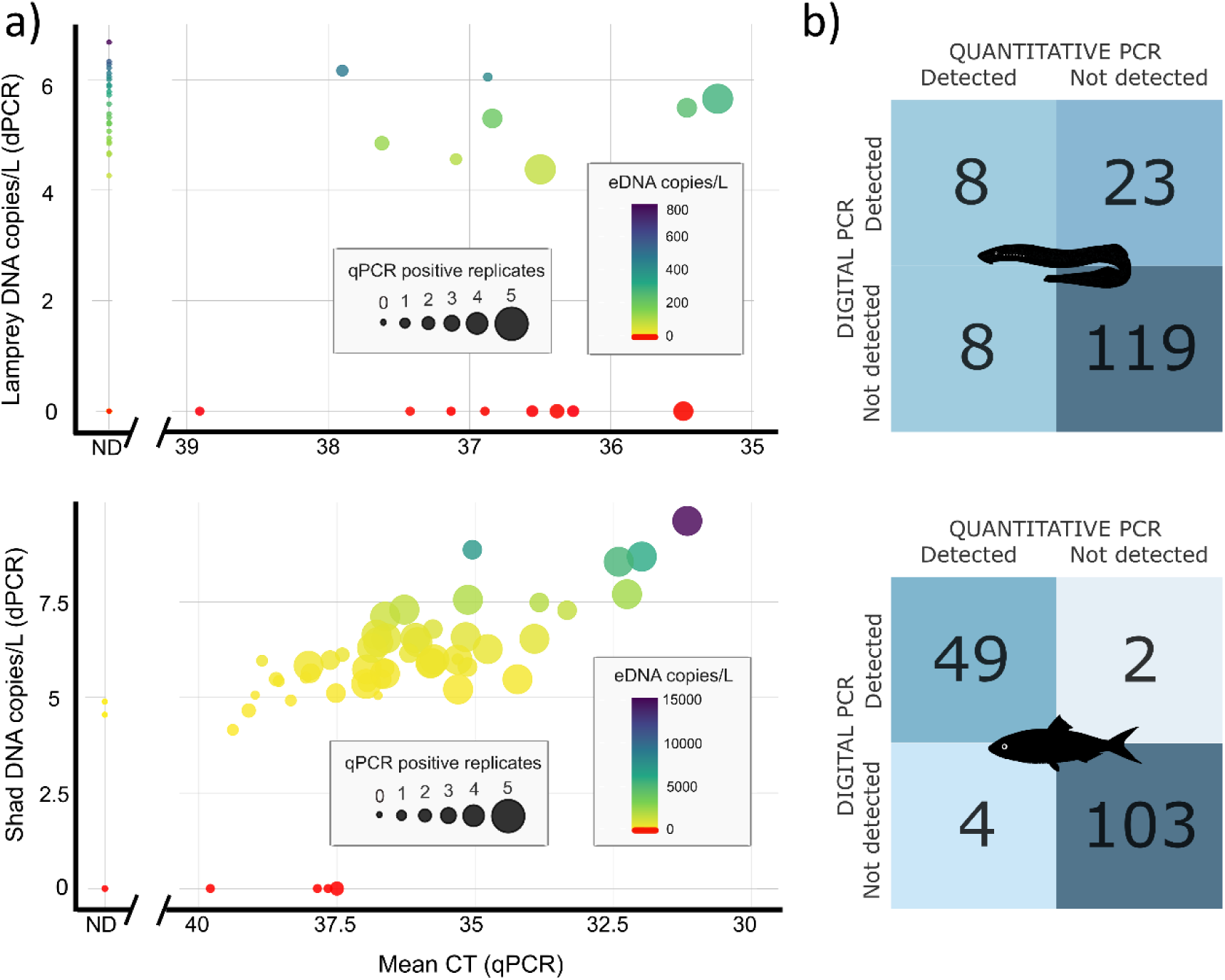
Comparison of qPCR and dPCR detectability outcomes for sea lamprey (top) and European shads (bottom). a) Correlation between CT values from qPCR and copies/L from dPCR. The bubble sizes are proportional to number of positive qPCR detections among replicates (n = 5) and colour represents the number of DNA copies per litre (in logarithmic scale). Each data point is based on five replications. ND in the x axis indicates no detection in qPCR. b) Confusion matrices for the sensitivity of the qPCR and dPCR approaches.

### eDNA abundance patterns along river stretches

We analysed the distribution patterns of sea lamprey and shads eDNA along river stretches, including four estuaries, using dPCR assays. Replicate samples from each sampling site were pooled to ensure a stable analysis. Statistical analysis revealed no significant differences between replicates for either sea lamprey (p-value = 0.9428) or shad (p-value = 0.3887), confirming the consistency of our sampling methodology. Overall, eDNA abundances of sea lamprey were significantly lower than those of shads (**Figure 5**). In the Basque Country, sea lamprey eDNA was mainly detected in the Bidasoa, Oiartzun, Urumea, Oria, Barbadun and Kadagua rivers. In Galicia, positive detections were recorded in the Eo, Anllóns, Umia, Ulla and Tambre rivers (**Figure 5a**). On the other hand, shads displayed a more restricted distribution, with detections confined to the downstream of the Bidasoa and Deba rivers in both years, while no detections were recorded in the estuary of any of the basins in 2019 (**Figure 5b**). The Bidasoa and Deba rivers were specifically analysed because all sampling points within these rivers showed positive eDNA detections. A consistent pattern of eDNA detection was observed, with higher concentrations found at downstream sites and progressively lower concentrations upstream, regardless of the sampling period (**Figure 5**).

**Figure 5.**
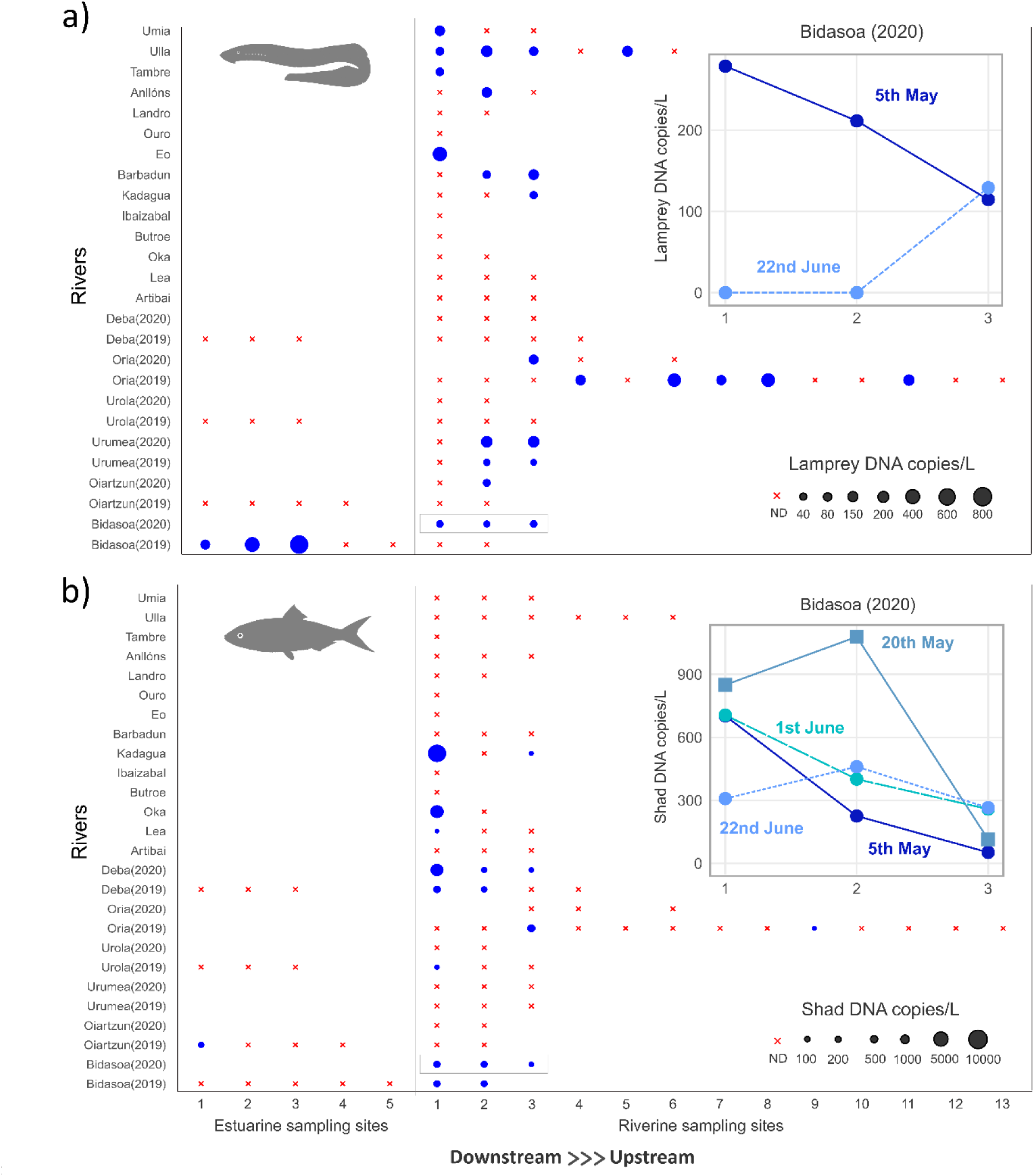
Spatial abundance patterns of eDNA detections using dPCR across Spanish sampling sites, where the dot size represents the eDNA concentration (copies/L) for a) lamprey and b) shads in each sampling point and red crosses indicate samples where no DNA was detected (ND). For Bidasoa river in 2020, the dots represent the average concentration (mean) of the five sampling dates within the year in which water was collected. Accordingly, the connected scatterplots show a breakdown of the results for the sampling dates within the year. Only results for those dates for which three sampling points were sampled and had at least one of them positive are shown.

## Discussion

In this study, we have built on existing knowledge of sea lamprey and European shad through a comprehensive research initiative spanning 44 river systems across Europe. This effort was made possible through international collaboration among researchers, enabling a large-scale application of the eDNA approach to assess the distribution of two diadromous species, both of significant conservation concern in the Northeast Atlantic region. Here we share key practical lessons from this ambitious endeavour, with the goal of guiding similar applications of eDNA for monitoring diadromous fish in similar and other contexts.

### Detection discrepancies among species

Here, we concurrently monitored the presence of both sea lamprey and European shad using a single eDNA sample. Both species are anadromous and exhibit similar habitat preferences during their spawning migrations. During spring, adult shad and sea lamprey migrate to rivers to spawn, after which most die (Silva et al., 2013). After hatching, juvenile shad then migrate to the sea in late summer and autumn (Aprahamian et al., 2003), while sea lamprey larvae (ammocoetes) remain buried in the riverbed for years before becoming parasitic juveniles (OSPAR Commission, 2009b). This buried larval stage, coupled with its generally declining population density (Almeida et al., 2023; Limburg & Waldman, 2009), might result in low eDNA detection rates. Conversely, shad, with a primarily pelagic habitat and high mobility during spawning, are more likely to release eDNA into the water column during migrations. According to these predictions, in our study, eDNA detection rates between the two species differed: shad was detected in 40% of the sites where it was expected whereas sea lamprey in 29%. These results emphasize the need to integrate species-specific behaviour patterns and habitat preferences in eDNA sampling design and result interpretation (Baltazar-Soares et al., 2022; Clemens et al., 2022).

### When and how to sample

The opportunistic nature of this international monitoring survey of diadromous fish led to distinct sampling approaches by each institution, influencing detection rates. These variations included sampling times, locations, collected water volumes, and filter pore sizes and types. Efforts were made to synchronize most of the eDNA sampling with the migration period of both species, assuming higher eDNA shedding rates during this time (Thalinger et al., 2019). However, this synchronization was not fully achieved for sea lamprey (Kelly & King, 2001; McIntyre et al., 2007; Silva et al., 2019; Silva et al., 2013; Taverny & Elie, 2009). Only in the Basque Country sampling was performed during upstream and downstream migration, while all other regions were sampled only during downstream migration. Both shad and sea lamprey eDNA were detected during upstream migration at expected locations, but sea lamprey was largely undetected, likely due to mismatches in timing with peak downstream migration and lower eDNA shedding rates in surface water. In contrast, the successful detection of sea lamprey was reported in Galicia as well as in the Tamar and Dee rivers, likely due to the high population density in these areas, which increased the likelihood of eDNA presence in the water samples. Interestingly, the detection rates did not appear to be significantly influenced by the filter pore size or type used, despite variations ranging from 0.22 μm to 1.50 μm. Larger pore sizes enhance the ratio of macro- to micro-organismal DNA, increasing the likelihood of detecting fish species (Deiner et al., 2018; Liu et al., 2024). Additionally, larger pore sizes facilitate the filtration of larger water volumes (Mächler et al., 2016), resulting in greater DNA yields. Despite using same pore size (0.45 μm), French samples, which used 30L of water volume, resulted in lower eDNA detection rates than Spanish samples, which used 1-2L of water volume. This suggests that smaller volumes of water do not necessarily lead to reduced detection. Similarly, we do not see a clear increase of detection rates when large pore sizes are used. Altogether, our findings indicate that when enough eDNA from a species is present, it can be detected regardless of the sampled water volume filtered or pore size used. However, given the low abundance of diadromous species in certain rivers, in particular beyond the upstream migration period, and in estuaries, where their concentration is even lower, using the largest water volume possible and a pore size larger than 0.45 μm is recommended to maximize eDNA based detectability.

### Methodological differences

The comparison between qPCR and dPCR derived results revealed differences depending on the species under study. For shad, results of both approaches are similar with 96% of the samples resulting in concordant results. In this case, only 6 out of 158 samples exhibited inconsistent results. The 4 qPCR positives with negative detections based on dPCR could be considered false positives, as they were characterized by only 1 or 2 positive replicates and high cycle threshold (Ct) values. Conversely, the 2 dPCR positive results that remain undetected in qPCR could be considered false negatives due to low DNA quantity, non-detectable by qPCR. For sea lamprey, the concordance rate dropped to 80%, with 31 out of 158 samples exhibiting differing results. Unexpectedly, 8 samples were found positive by qPCR but negative by dPCR. Of these, 6 samples had only one or two positive qPCR replicates; however, 2 samples had three and four positive replicates with lower Ct values, indicating potential dPCR false negatives. dPCR utilizes extensive sample dilutions divided into thousands of independent partitions assessed through Poisson statistics (Qiagen, 2024). Significant dilution can produce partitions that do not contain the target DNA, especially at low initial concentrations (Whale et al., 2013), which might explain the observed false negatives. Additionally, 23 samples that were positive in dPCR but negative in qPCR highlight potential limitations in the sensitivity of the qPCR method for detecting low concentrations of DNA. Overall, dPCR demonstrates superior efficacy compared to qPCR for detecting species within low-abundance populations. dPCR is also advantageous as it provides precise quantification of number of eDNA copies, which allows absolute abundance comparisons between samples. Yet, to ensure reliable dPCR results, sample concentrations must be maintained within the dynamic range to reduce partitioning errors, particularly for unknown concentration (Qiagen, 2024). Systematic optimisation of dilution factors improves the confidence of dPCR in identifying low target DNA concentrations, thereby reducing false negative rates. This optimization process involves adjusting the dilution levels to achieve the desired sensitivity and precision, which is particularly important when dealing with low-abundance targets (Jiang et al., 2022). While in general dPCR excels in sensitivity and accuracy, some applications might prefer using qPCR due to its lower cost, important particularly for large-scale monitoring (Zhang et al., 2024). This trade-off between cost and accuracy highlights the importance of selecting the appropriate method based on specific research objectives and available resources.

### Can eDNA be used as reliable sources of presence/absence for diadromous fish?

The finding from this survey revealed inconsistencies between eDNA detection and historic expected occurrences of sea lamprey and shad. Focusing on sites where the species were expected, we identified false negatives, which could be attributed to low abundance of the species, resulting in low eDNA concentrations in the dPCR or qPCR assays. Because false negatives can misrepresent species presence and hinder conservation efforts (Desrochers et al., 2010), it is important to reduce them by adapting sampling times to coincide with peak migration, especially during spawning periods, using larger pore size filters and filtering large volumes of water, and applying more sensitive methods such as dPCR. Focusing on the sites where the species was not present, only in the case of shad we detected 3 apparent false positives; however, one of them occurs only in the qPCR results, in one replicate sample of two, in one out of 5 qPCR replicates and with high Ct values (39); the other two positive locations (Inny and Ilen rivers) did not have dPCR results available, but the qPCR positives were based on a single replicate out of 5 and with high Ct values (37 & 39). In view of this, all these three apparently false positives can be considered true negatives instead. Focusing on the sites for which presence or absence in the species was unknown, all cases resulted negative for lamprey. However, for shad, we revealed new sites where the species could be present. Interestingly, in river Deba the initial reference database was marked as “unknown”, but was changed to “expected” when, after seeing the eDNA results, the monitoring agency returned to the river and did find shad (Ekolur, personal communication). For this study, the availability of a comprehensive reference database has enhanced our understanding of the eDNA based results, facilitated interpretation and identification of potential causes of false negatives and positives. Therefore, we propose eDNA as a complementary information source for diadromous species monitoring, which can be applied to increase time and space coverage due to its easy logistic, non-invasiveness and cost-effectiveness.

## Supporting information

resulting in a total of 313 samples (Figure 1; Table S1). The

unknown and 38 not expected sites (Figure 2a; Table S3).

sample filtered ranged from 1 to 30 litres (Table S2). Spanish

## Acknowledgements

This work has been funded by the INTERREG Atlantic Area program through project “DiadES - Assessing and enhancing ecosystem services provided by diadromous fish in a climate change context” (EAPA_18/2018), by the Department of Economic Development and Infrastructure of the Basque Government through project “GENGES - Development and application of genetic methods to improve marine ecosystem management”, by the Spanish Ministry of Science and Innovation through project “EDAMAME - Environmental DNA based approaches for marine and aquatic monitoring and evaluation” (CTM2017-89500-R), by the Game & Wildlife Conservation Trust, Queen Mary University (QMUL) of London and Cefas’ Science Excellence fund. This work was performed under at doctoral grant awarded to Mukesh Bhendarkar by the Indian Council of Agricultural Research (ICAR), Government of India. The authors would like to thank Oriol Canals for his valuable suggestions during the initial stage of the project, Michał Skóra and Hui Wei for their help with sample collection, as well as Environment Agency, Natural Resources Wales, Game and Wildlife Conservation Trust, Marine Scotland Science, Scottish Fisheries Co-ordination Center, Ness District Salmon Fishery Board and Dr Miran Aprahamian for providing data, as well as expert opinion on sea lamprey and shads presence at sampled sites in the UK. Finally, we would like to thank Gordon H. Copp for the expert advice that he contributed to this work before he sadly passed away in 2023.

