## Supplementary material for "Lessons learned from applying eDNA surveying to diadromous fish detection across the north-east Atlantic region": sample filtered ranged from 1 to 30 litres (Table S2). Spanish

**Table S2** Overview of eDNA sampling and processing of collected samples across the different regions indicating the year, period, number of samples and replicates, and other sampling features. Field control corresponds to negative control where ultra-pure water filtered alongside collected field samples, to assess for possible contamination leading to false positives. In all cases PCR controls were included.

| **Region** | **Sampling year** | **Sampling months** | **Number of samples** | **Number of replicates** | **Field Control** | **Volume of water sampled** | **Filter pore size** | **Filtration** | **Extraction method** |
| --- | --- | --- | --- | --- | --- | --- | --- | --- | --- |
| Basque (Spain) | 2019-20 | 5,6,10 | 161 | 2 | Yes | 2 L | 0.45 μm | Field | Qiagen QIAamp DNA Kit |
| Galicia (Spain) | 2020 | 9 | 22 | No | No | 1-2 L | 0.45 μm | Lab | Qiagen QIAamp DNA Kit |
| France | 2021 | 7,8,9 | 9 | 2 | No | 30 L | 0.45 μm | Field | Machery-Nagel Düren NucleoSpin®Soil kit |
| Ireland | 2019 | 8,9,10,11 | 68 | 3 | Yes | 2L | 1.50 μm | Lab | Qiagen QIAamp DNA Kit |
| United Kingdom | 2019 | 8,10 | 53 | 3 | Yes | 3 L | 0.22 μm | Field | DNeasy PowerWater Sterivex Kit |
